# BioEncoder: a metric learning toolkit for comparative organismal biology

**DOI:** 10.1101/2024.04.03.587987

**Authors:** Moritz D. Lürig, Emanuela Di Martino, Arthur Porto

## Abstract

In the realm of biological image analysis, deep learning (DL) has been a transformative force. Tasks like finding areas of interest within an image (segmentation) and/or sorting images into groups (classification), can now be automated and achieve unprecedented efficiency and accuracy. However, conventional DL methods are challenged by large-scale biodiversity datasets, which are characterized by unbalanced classes and small phenotypic differences between them. Here we present BioEncoder, a user-friendly toolkit for metric learning, which overcomes the aforementioned challenges by focussing on learning relationships between individual data points rather than on the separability of classes. This approach simplifies training of robust image classification models for biologists, democratizing access to advanced DL techniques. BioEncoder is encapsulated in a Python package, crafted for ease of use and flexibility across diverse datasets. It features taxon-agnostic data loaders, custom augmentation options, and streamlined hyperparameter adjustments through text-based configuration files. The toolkit’s significance lies in its potential to unlock new possibilities in biological image analysis - from phenomics and disease diagnosis to species identification and ecosystem monitoring. BioEncoder focuses on the urgent need for toolkits bridging the gap between complex DL methodologies and practical applications in biological research.

## Introduction

The advent of advanced digital imaging technologies has revolutionized biological research, particularly in the realms of medicine, agriculture, biology, and notably, within the vast collections of natural history museums (NHM) (Blackburn et al., 2024). These technologies have enabled the creation of high-quality digital images of biological specimens at unprecedented rates, capturing intricate details of organism structure, function, and behavior (Johnson et al., 2023; Nelson & Ellis, 2018). Images have become a cornerstone in understanding the rich biodiversity encompassed in these collections, but the sheer volume and complexity of this digital imagery presents additional challenges: as the reservoir of digital images expands, so does the need for sophisticated methods to analyze and interpret these data (Johnson et al., 2023; Lürig et al., 2021). A central challenge in biological image analysis lies in extracting meaningful features from high-dimensional images that effectively represent the underlying biological structures or differences among specimens. This task becomes increasingly difficult when dealing with the diversity of specimens typical of natural history collections (Van Horn et al., 2021). In these collections, rare species and high sampling efforts often result in datasets with unbalanced classes, encompassing specimens that exhibit only subtle phenotypic differences and complex hierarchical taxonomic relationships (Belitz et al., 2022). Such intricacies necessitate advanced analysis methods capable of accurately capturing and interpreting these subtle differences.

In response to these challenges, recent advancements in deep learning (DL) have been pivotal. DL has the capacity to automate image segmentation and classification, and thereby make image analysis more efficient, accurate, and objective (Høye et al., 2021; LeCun et al., 2015; Lürig et al., 2021). Specifically, DL models have been used in tasks such as identifying and categorizing species, detecting morphological differences, and analyzing patterns of development and disease (McQuin et al., 2018). By automatically defining and learning features from images with high accuracy and throughput, DL allows for a more comprehensive and high-throughput understanding of biological images. The automation of these tasks not only streamlines the workflow but also opens the door to new possibilities in biological research, such as large-scale phenotypic screening (Jackson et al., 2019), automated classification for biodiversity assessment (Van Horn et al., 2018), and detailed morphological analyses (Di Martino et al., 2023) that would be impractical, if not impossible, to perform manually. However, traditional DL approaches are hampered by their inability to handle unbalanced classes effectively and their reliance on non-metric feature spaces, which are not conducive to quantitative comparisons between classes. Metric Learning (MEL) emerges as a variant of DL, distinguished by its ability to learn features of input images on a metric space (Bellet et al., 2022). This space enables a quantitative comparison of specimens or groups according to a chosen distance metric (e.g. Euclidean) in a high-dimensional feature space. MEL approaches are often used in person re-identification, image retrieval, and other visual recognition tasks with extreme numbers of classes and heavy-tailed distributions (Song et al., 2015; Xun Yang et al., 2018), as it focuses on relationships between individual data points rather than on the separability of classes (Smith et al., 2023). In biological image analysis, this approach is crucial as it allows the model to capture and emphasize important features across diverse classes, including those that are underrepresented or exhibit subtle phenotypic differences, such as those due to seasonality, among others (e.g. Belitz et al., 2022). MEL thus facilitates a more quantitative and accurate representation of the vast and varied biological specimens found in natural history collections.

Despite the potential of MEL, its integration into biological data analysis pipelines has been slow, primarily due to the lack of user-friendly toolkits that incorporate recent advances in this field. To bridge this gap, we introduce BioEncoder, a comprehensive toolset designed to facilitate biological image classification and the learning of features from images through metric learning (Fig. 1). BioEncoder is tailored for taxon-agnostic analysis, making it applicable to any species, and particularly suited to the diverse datasets typically found in natural history museums. The package provides an intuitive and flexible implementation, featuring data loaders, custom augmentation techniques, and hyperparameter selection through YAML configuration files, thus allowing for easy customization and application to various datasets and research questions. Our evaluation of BioEncoder’s performance on a biological image dataset of the polymorphic damselfly species *Ischnura elegans* demonstrates its efficacy (Le Rouzic et al., 2015; Willink et al., 2019, 2023). The package includes rich and interactive model visualizations, enabling detailed analysis and interpretation of the learned features. This aspect is crucial in understanding phenotypic differences captured in the images, and thereby the nuances of biodiversity. In summary, the BioEncoder package has the potential to improve applications like disease diagnosis, species identification, and morphometrics, which marks it as a valuable tool in the broader field of organismal biology, and especially within the context of natural history collections.

**Figure 1.**
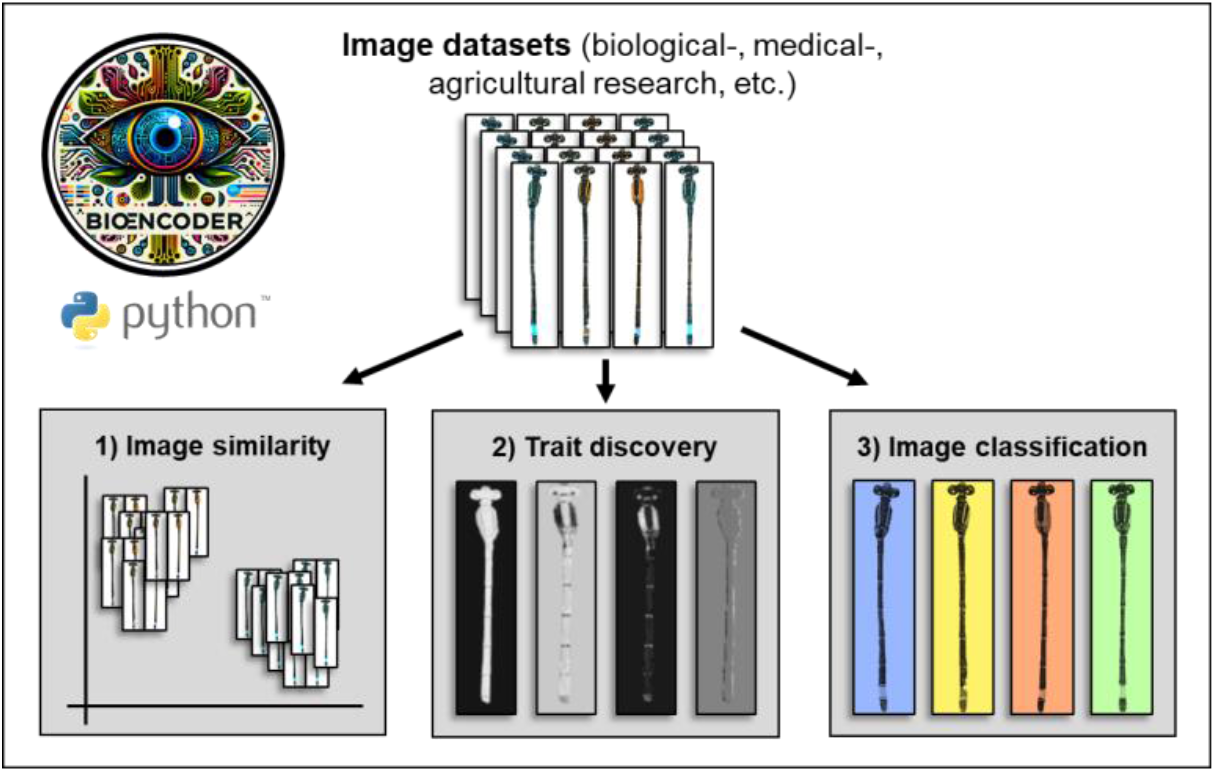
BioEncoder is a Python package for Metric Learning; providing functionality to 1) quantify image similarity, 2) to discover traits of relevance, and 3) classify images. The package’s focus lies on user friendliness, and it can be used in interactive Python sessions or through entry points from the command line.

## Materials and Methods

### The package

BioEncoder is a Python package that provides user-friendly access to advanced and scalable MEL techniques that can be flexibly applied without deep computational knowledge. Although we conceived the package with applications in evolutionary ecology and organismal ecology in mind, its utility can be applied to image datasets from any scientific domain (e.g., biomedical or agricultural research). The three main application contexts of the package are (Fig. 1):

1. image similarity: compare two images and calculate a similarity score
2. trait discovery: identify areas that are important for classification
3. image classification: predict image class based on image features

BioEncoder was designed to be user-friendly, and its entire functionality is wrapped into a total of nine public functions (Fig. 2), which simplifies and facilitates user interaction. The current version (0.1.1, snapshot available through the data-repository (Lürig et al., 2024)) is implemented in Python 3.9, and contains most of the state-of-the-art metric losses, such as Supervised Contrastive (SupCon)(Khosla et al., 2020), Additive Angular Margin Loss (ArcFace) and Sub-center ArcFace losses (Deng et al., 2020), as well as more traditional metric losses, such as triplet loss. Major dependencies include *pytorch(Paszke et al., 2019), pytorch-metric-learning(Musgrave et al., 2020)*, and *timm(Wightman, 2019)*. We tested the package on Windows and Unix systems.

**Figure 2.**
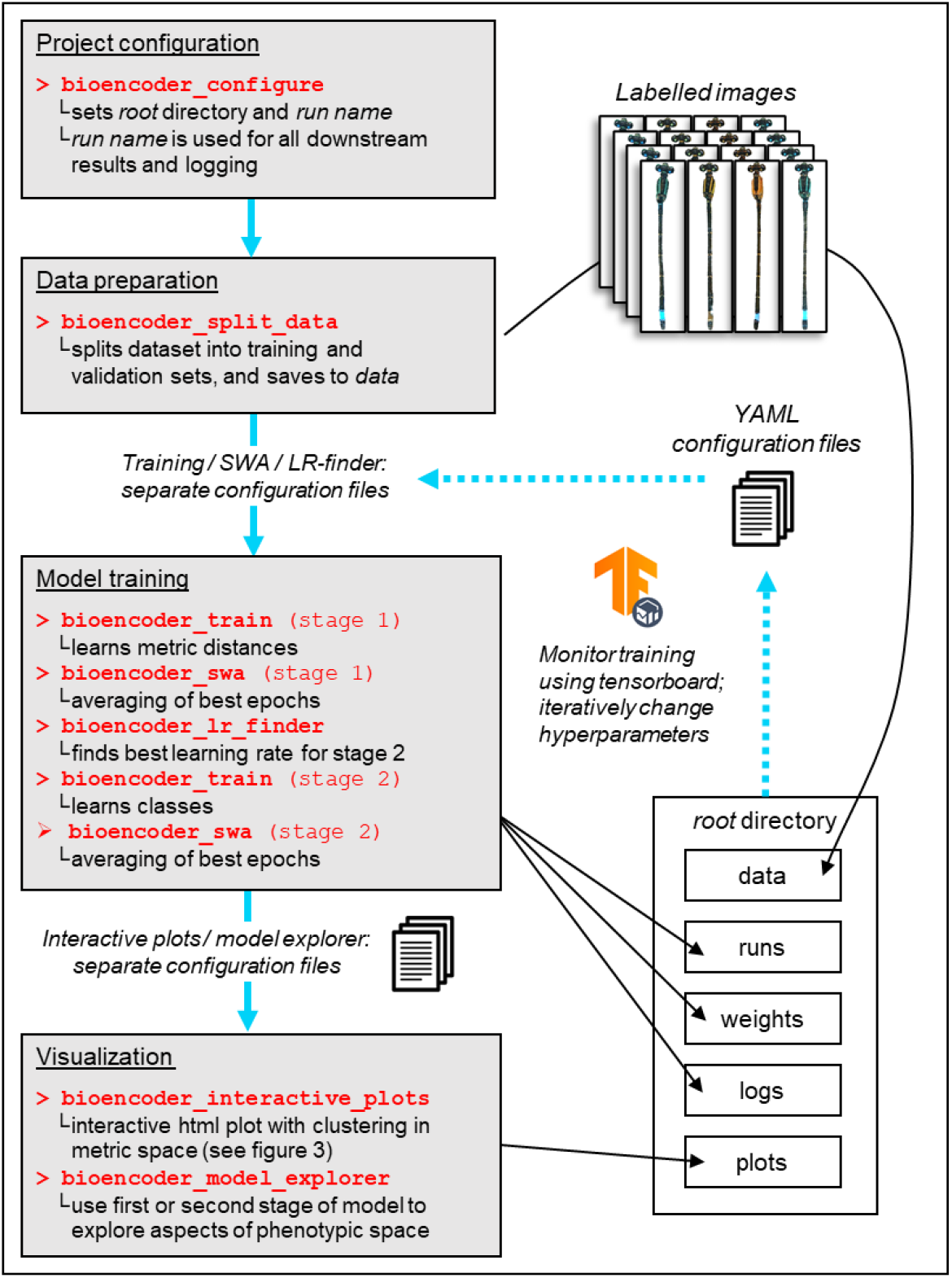
The BioEncoder workflow. Initially, users specify an existing *root directory*, where all materials and results that are produced in the training process are stored within subdirectories, and a *run name*, which is added as a prefix to all data, weights, logs, etc. that are stored within the subdirectories. After splitting the data and adding it to the data directory within *root*, model training can be started with a suitable configuration file in YAML format (templates are provided through the data repository (Lürig et al. 2024).

### Installation and quickstart

BioEncoder can be used either through a command line interface (CLI - .e.g, the windows terminal or the linux console), or interactively through an integrated development environment (IDE - e.g., VS code or Spyder). In either case, the package is installed from the Python Package index using ‘pip install bioencoder’. We highly recommend installing the package and all necessary dependencies to a Python virtual environment. In short; to test the package’s functionality, users should i) install a suitable python package and environment manager, such as conda or mamba, ii) create an environment with a Python 3.9 kernel, iii) install the package and its dependencies, iv) download the example image dataset we provide (Lürig et al., 2024), and v) follow the instructions in the repository. More detailed instructions for how to get started can be found on the package’s github repository (https://github.com/agporto/BioEncoder).

### Case study

In the following we demonstrate the functionality and performance of BioEncoder using a biological dataset of the polymorphic insect *Ischnura elegans* (Common Bluetail, Fig. 3A), which exhibits some typical properties of biological datasets: highly unbalanced classes and phenotypic gradualism. *I. elegans* is a short-lived univoltine damselfly, which expresses three female color morphs after emergence (androchrome, infuscans, and infuscans-obsoleta [hereafter, A-, I- and O-morph], Fig. 3B). The A-morph is thought to mimic the monomorphic males to avoid negative fitness consequences stimming from male mating harassment (Le Rouzic et al., 2015). The Svensson group maintains a long term monitoring program to study frequencies and evolutionary dynamics of the morphs in multiple populations close to the town of Lund (Skane, Sweden). The damselflies are caught with hand nets and brought back to the laboratory, where morphs are identified and imaged under highly standardized conditions (for details see (Le Rouzic et al., 2015)). In the monitored populations sex ratios and morph frequencies are heavily skewed (on average: males=60%, A-morph=26%, I-morph=12%, O-morph=2%)(Le Rouzic et al., 2015). The gradual phenotypic differences due to color polymorphism and male mimicry, as well as the strongly imbalanced groups make this dataset highly suitable for a case study using BioEncoder. All of the preprocessed images, configuration files and the script used for training are available through a data repository (Lürig et al., 2024). The following paragraphs match the workflow presented in Fig. 2 - additional details can be found in the Supplementary Information (SI).

**Figure 3.**
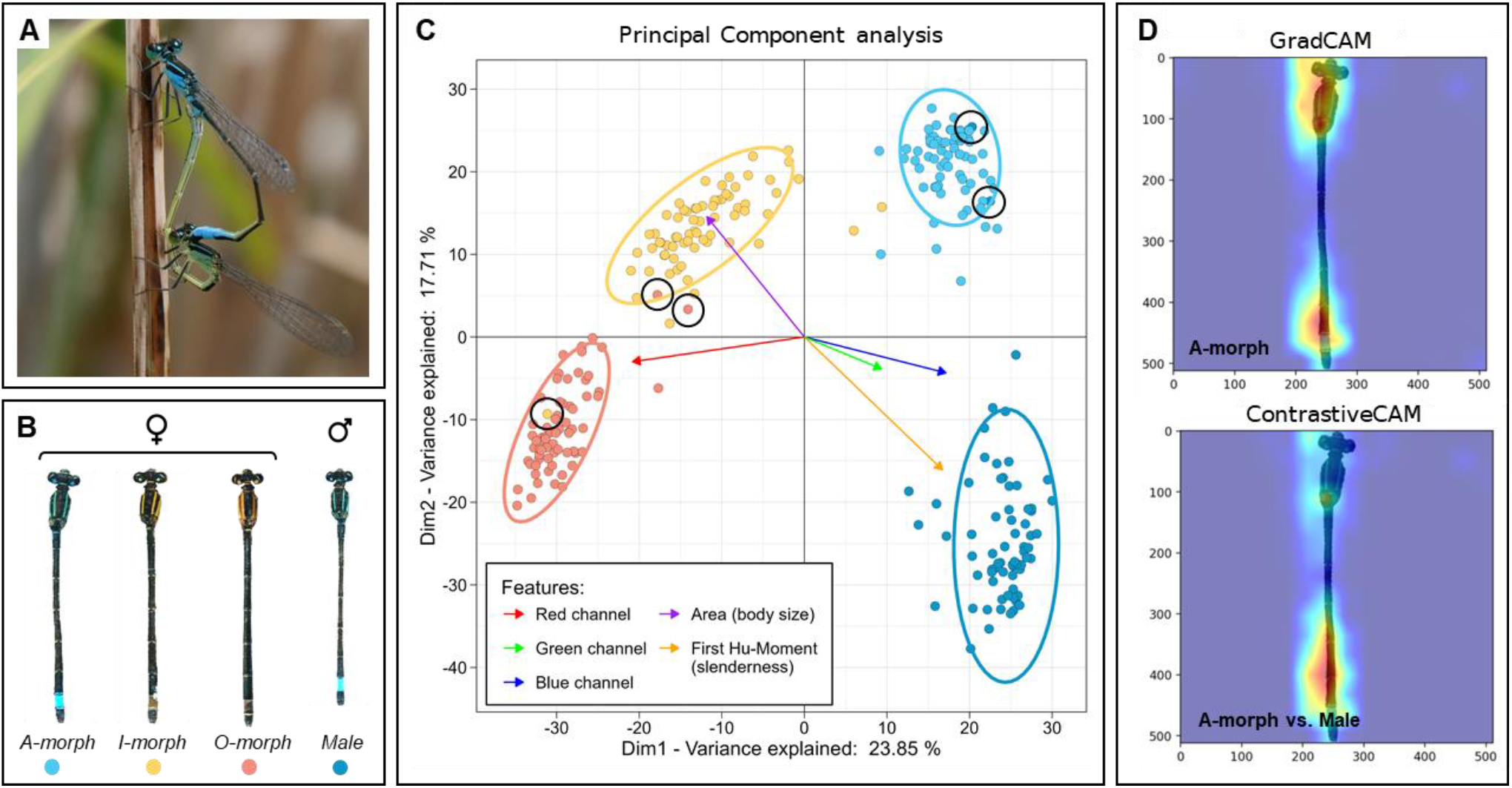
BioEncoder case study. **A:** The damselfly *Ischnura elegans* has a widespread distribution across Europe and parts of Asia. We used a dataset of 8386 standardized images for the case study. **B:** *I. elegans* expresses three female morphs, one of which (A-morph) is a putative mimic to the monomorphic male. **C:** Principal Component Analysis of the image embeddings (length = 2048) of the validation set (n=70 per class), extracted using the three top performing checkpoints from the first stage. Each point is a single damselfly, color coded by morph ID (black circles indicate falsely labeled images). The colored arrows indicate the loadings for five quantitative support variables, which we collected from the images using the phenopype package to inform the embedding space: mean pixel intensity in the three color channels (red, green, blue), as well as absolute area in pixels, and the first Hu-moment (measure for slenderness). While the three female morphs separate mostly by color and color patterning, the A-morph is, as expected, intermediate to I- and O-morph and males along a gradient of size and shape. BioEncoder is able to generate interactive ordination plots, so that when a user hovers over a data point they will see a pictogram of that specific individual. **D:** Class activations and areas of relative importance for class discrimination; extracted using the final checkpoint from the second stage. GradCAM shows that for the A-morph, the head, thorax and lower abdomen region are highly informative. When compared to the male using ContrastiveCAM, the model uses information on abdomen slenderness, which appears important given that it would be very difficult to differentiate the putative mimic on coloration alone.

#### Configuration

We initiated the workflow using BioEncoder’s main configuration function, which is used to point to a root directory where all output is saved, separated by different run-names. After initial setup, we supplied separate YAML configuration files for the two stages of the training process, as well as for model averaging, visualization and the model explorer. In each YAML file users need to specify the model architecture, hyperparameters, and augmentations. The configuration files used for the case study can be found in the data repository (Lürig et al., 2024) as well as the github repository (https://github.com/agporto/BioEncoder), and may serve as a starting point for different data sets and use-cases.

#### Data preparation

The dataset we used spans a period of 10 years (2014 - 2023), of which we used 6 years for training and validation, and the remaining 4 years for final testing. In total, the training and validation set contained 8386 images (5000 males, 2240 A-morph, 999 I-morph, 147 O-morph). When splitting the data into training and validation sets, we allowed for a maximum spread of 6 times the smallest class (6*147 = 882), which left us with 2793 images (882 males, 882 A-morph, 882 I-morph, 147 O-morph). Users can set their own maximum ratio between classes to reduce the weight of abundant classes and train a more generalizable model. We used BioEncoder’s default “flat” option for the train/validate split, which uses 10% of all images for validation; evenly distributed across all four classes to ensure that validation metrics are not influenced by the dominant classes (276 images = 69 per class). However, other split options are available and can be chosen by users to accommodate their specific data set or training context.

#### Model training (stage 1)

We trained the model using a 64-core workstation with an RTX A600 GPU (48GB of GPU memory), CUDA 12.1, and cuDNN 8.9.2. For the first stage training, we employed a pre-trained EfficientNet-B5 model, chosen for its proven efficacy in feature extraction due to pre-training on a diverse dataset (JFT, (Tan & Le, 2019)). This stage was trained for 100 epochs, but did not incorporate a classification head, reflecting the focus on representation learning over direct class prediction and aligning with a supervised contrastive learning approach. We used the SupCon loss function to optimize the embedding space by reducing intra-class variance and increasing inter-class separation, which was instrumental in enhancing the model’s ability to distinguish between distinct instances. Images were resized to 384x384 pixels, and augmented to introduce variability into the training data, thus aiding the model in learning robust and invariant features (for details on training parameters and augmentations see SI).

#### Model training (stage 2)

Upon completion of the first stage of training, we employed stochastic weight averaging (SWA) on the top three performing model weights to further enhance the generalization capabilities of the EfficientNet-B5 model. Subsequently, to optimally configure the model for the second stage of training, we used the package’s learning rate finder script to empirically determine the most effective learning rate. This method is based on a short training where the learning rate is incrementally adjusted, thereby identifying an optimal rate that balances training speed and convergence stability. The second stage was trained for 30 epochs and introduced a classification head to the previously frozen EfficientNet-B5 architecture, transitioning the model’s focus from representation learning to direct class prediction. Note that the second stage of training is completely optional and does not typically result in increased classification accuracy. Upon completion of the second stage of training, we employed SWA on the top three performing model weights to retrieve the final model.

#### Visualization

We created an interactive plot using the SWA weights of the first stage (Fig. 3C). The panels show ordinations of the image embeddings for each individual from the validation set using Principal Component Analysis (PCA); t-distributed Stochastic Neighbor Embedding (t-SNE) is also available. The PCA shows the separation of the individuals and their clustering in linear space. Moreover, we used the model explorer functionality of BioEncoder on the SWA weights of the first stage to visualize the filters, saliency- and activation-maps of specific layers from the training process (Fig. 4). Specifically, we explored class activation maps to identify the parts of an input image that most impact the final classification (GradCAM), and to identify parts of an image that differ most from another class (ContrastiveCAM).

**Figure 4.**
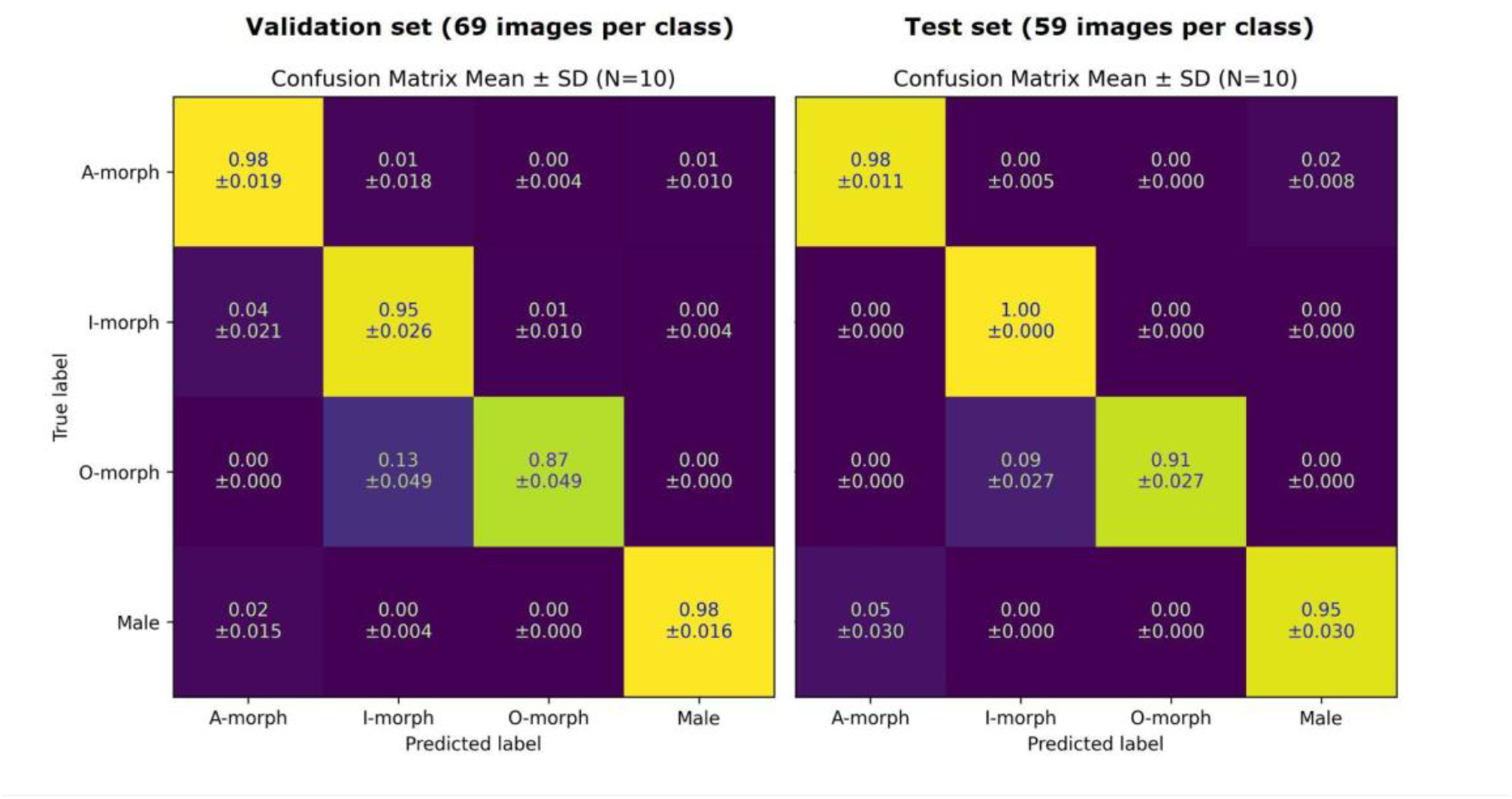
Confusion Matrix for validation (N=69) and test set (N=59). In both data sets, classification accuracy was high (95% - 100%) for A-morph, I-morph and males, whereas accuracy for the O-morph was somewhat lower (87% in the validation set, 91% in the test set). The O-morph was most often confused with the I-morph, which, aside from the lower amount of training images, may also be due to the presence of several putatively mislabelled individuals (Fig. 3C).

We repeated the entire workflow 10 times, including all steps of the training process, SWA, and learning rate finders. The only difference between the runs was the random seed for the selection of the training and validation images. We present accuracy and confusion scores for validation and test sets below (Fig. 4).

## Results

The first stage of the BioEncoder pipeline trained for 100 epochs, which took on average 81.273 ± 0.199 minutes (mean and standard deviation for the 10 runs) on the test system, with significant improvements in both the training and validation loss metrics (see SI for details). The encoder validation metrics showed a consistent increase in precision at one, from its initial value of 0.564 ± 0.056 to an average of 0.94 ± 0.012 by the end of the training period. In the second stage of the pipeline, model training extended over an additional 30 epochs (4.358 ± 0.0214 minutes). Accuracy tended to fluctuate around a high level without significant change, i.e., from an initial 0.946 ± 0.015% to 0.941 ± 0.017%, just like the F1 scores which ranged from 0.95 ± 0.012 at the beginning of the training period to 0.945 ± 0.015 by its final epoch. As the aggregated confusion matrix of the validation set indicates (Fig. 4), all classes were correctly predicted with 95 and 98% accuracy, except the O-morph with 87%, which was most often confused with the I-morph. Moreover, confusion scores of the validation sets (69 images per class) were similar to the scores of the test set (59 images per class), which the model had not seen until the final inference.

Using the normalized image embeddings we discovered two relevant axes of separation among damselfly morphs and sexes (Fig. 3C). On the one hand, all four groups separate along a gradient of coloration and patterning, which is largely aligned with the first dimension on the x-axis. Most notably, the O-morph with its uniform reddish thorax coloration is on the far left and clearly separated from the I-morph, which has a yellow-striped thorax. To the far right are A-morph and the males, which have a blue-striped thorax and a bright blue color patch at the end of their abdomen. On the other hand, female damselflies group together on one end of a size and shape gradient, with the larger, more stout O- and I-female morphs toward the upper left, the smaller, more slender males toward the lower right, and the A-morph, the putative male mimic, in between. The proximity of the A-morph to the males and its intermediate position between the other morphs and the males in phenotypic space provides strong quantitative support for the hypothesis of male-mimicry, but it also shows that the mimicry is more complete in terms of coloration and patterning than in body size and shape. Careful visual inspection of feature space also revealed that some images in our dataset were most likely incorrectly labeled (see black circles in Fig. 3C).

The classification layer of the second stage allowed us to explore class specific activation maps and discriminative features across classes. We found that regions with high informative value largely overlapped with the regions typically used for identifying the morphs by human experts. Specifically, inspections of the visual output from the GradCAM method revealed that all classes had high activations in the region around thorax and abdomen (Fig. 3D). In addition, we found high activations in the region of the head, which might be driven by the small color patches above the eyes and potentially head shape. ContrastiveCAM revealed strong phenotypic differences between morphs and sexes, such as thorax coloration (monochrome in O-morph vs. striped in A-/I-morph and males), lower abdomen coloration (blue patch present in A-morph and males vs. absent in I- and O-morph) and abdomen shape (slender in males vs. stout in females) (Fig. 3D). We used the upper layers of the model for both GradCAM and ContrastiveCAM, which provide low resolution but provide a good overview. More fine scale variation could be explored by focussing on the lower layers of the model.

## Discussion

The classification of specimens and species is one of the most pervasive image analysis tasks in organismal ecology and evolution (Høye et al., 2021; LeCun et al., 2015; Lürig et al., 2021). However, biological datasets pose great challenges to traditional DL approaches (Van Horn et al., 2021), and quantitative comparisons between classes or relating any learned features to underlying biological structures has remained unfeasible. With BioEncoder, we present an easy-to-use toolbox that uses MEL to encode image data into a robust metric feature space, and to classify images with high accuracy (Bellet et al., 2022). In developing BioEncoder, our primary aim was to bridge a significant gap in the accessibility of advanced metric learning tools for the biological community, rather than redefining the cutting edge of DL for image classification. Recognizing the unique challenges that biologists face with the often unbalanced data distributions inherent in natural history and organismal biology, we tailored BioEncoder to serve as an intuitive, user-friendly toolkit. By prioritizing ease of use, we strive to democratize the application of MEL, a methodology we believe is uniquely suited to biological research. In particular, MEL’s focus on understanding the relationships between individual data points, rather than merely the separability of classes (Song et al., 2015; Xun Yang et al., 2018), aligns closely with the granularity and diversity of biological data.

Through BioEncoder, we also aim to enhance our ability to derive insights and/or interpretation from image classification models. The toolkit is designed to offer comprehensive model visualizations, alongside interactive PCA and t-SNE plots, specifically tailored to facilitate biologists in navigating the embedding spaces generated by the models (van der Maaten, 2008). This feature is instrumental in translating the complex, high-dimensional embedding space into a more intuitive framework of traits, mirroring the way biologists traditionally encode and interpret imaged data. For instance, the feature space that we constructed by extracting image embeddings from the damselfly dataset confirms several hypotheses and observations from field observations (Le Rouzic et al., 2015) and experimental research. Firstly, female morphs form distinct phenotypic clusters along a gradient of body coloration and patterning, as indicated by the clear separation along Dimension 1 in the PCA (Fig. 3C). Moreover, the A-morph cluster is located between the males and I- and O-morph along a gradient of size and shape, supporting the observation that A-morph females are somewhat smaller and more slender than the two other morphs. This observation, as well as the male-like blue thorax and abdomen coloration, provides quantitative support for the existing hypothesis of male mimicry in the A-morph (Le Rouzic et al., 2015; Willink et al., 2019, 2023).

In terms of pure robustness and performance, we observe high classification accuracy for all morphs, despite the presence of falsely labeled images (Fig. 3C), and including the O-morph, for which only a fraction of the images used in the other classes were available. This suggests that our training setup provided an effective approach to learning discriminative features from the damselfly dataset in the first stage, leveraging the capabilities of the chosen architecture and the contrastive learning paradigm to prepare the model for subsequent stages (Smith et al., 2023). Such MEL approaches are often used in person re-identification, image retrieval, and other visual recognition tasks with extreme numbers of classes and heavy tailed distributions (Song et al., 2015; Xun Yang et al., 2018). In the damselfly dataset, the differences between morphs and sexes are very subtle, and the untrained eye would have difficulties discriminating, for example, the males from the A-morph (Le Rouzic et al., 2015). Such male mimicry is a special case of intra-specific phenotypic similarity, and there are many other cases of intra- and interspecific mimicry, which have repeatedly evolved (Ezray et al., 2019). In general, the tree of life forms a diverse network of species with varying degrees of relatedness and phenotypic similarity across its branches, which makes an extremely challenging case for classification models (Stevens et al., 2023). By applying BioEncoder to an extreme case of phenotypic similarity across classes, we show that the MEL approach is also effective when used on biological datasets.

Despite the promising results obtained from the experiments, there are still some limitations to BioEncoder that need to be addressed. For instance, the tool relies on manual tuning of hyperparameters, which can be time-consuming and requires expertise in DL (LeCun et al., 2015). Furthermore, the experiments conducted in this work were limited to a few datasets and taxa, and it remains to be seen how well the tool performs on more diverse datasets (Stevens et al., 2023). Still, the results obtained from the experiments demonstrate the effectiveness of the proposed approach in capturing biologically relevant features that can be used for classification tasks. This adaptability is further highlighted by BioEncoder’s capacity to encode images not previously seen in the training set, showcasing its robustness and potential for continuous learning. Such a feature is particularly valuable in addressing the ‘long tail’ problem of biodiversity, where the vast majority of species are rare and underrepresented in scientific datasets. BioEncoder offers a scalable solution to such challenges by efficiently encoding diverse and sparse phenotypic information. We hope that the user-friendliness of the package, as well as the provision of extensive support materials (Lürig et al., 2024) will encourage and support a wide range of users in their attempts to capture the full spectrum of biological diversity.

## Conclusion

In conclusion, BioEncoder presents a valuable tool for encoding biological images into meaningful subspaces, which can be used for various applications in organismal biology. The tool provides a taxon-agnostic approach that is applicable to any biological dataset, and the use of metric learning allows for the generation of features that are more biologically meaningful and transferable across different datasets. With further development, BioEncoder has the potential to advance the field of organismal biology and facilitate the discovery of new biological traits and relationships.

## Author contributions

AP, MDL and EDM conceived the idea. AP implemented the first draft of the package, and provided a partial draft of the manuscript. MDL provided the dataset for the case study, gave critical feedback on the package, implemented the pypi-version,and led the writing after the first draft. AP and EDM acquired funding. All authors critically contributed to the final draft of the paper.

## Acknowledgements

MDL was supported by a Marie Sklodowska Curie Individual fellowship awarded by the European Commission (PhenoDim; Grant No. 898932). AP was supported by a Hardware Grant from NVIDIA Inc. EDM was supported by the Research Council of Norway (Grant No. 314499). The damselfly dataset was collected by the Svensson lab at Lund University, Sweden, and all images as well as the labels were kindly provided by Erik Svensson.

## Conflict of interest

The authors declare that this research was conducted in the absence of any conflict of interest.

## Supplementary Information

### Materials and Methods

#### Preprocessing

Prior to training, we used a custom pipeline based on phenopype (Lürig, 2022) to standardize the specimens for better training performance and accuracy. Specifically, we i) segmented the damselfly body without wings and appendages, ii) rotated the segmented specimens so all of them are facing upwards, iii) mounted the segmented and rotated body onto a white canvas, and iv) resized the final image to 512x512; preserving relative size differences among specimens. Note that images are generally not required to have the same dimensions, as they are resized by BioEncoder during the training using the parameters defined within the YAML configuration files.

#### Augmentation

We applied the following augmentations through the configuration files: RandomResizedCrop, Flip, RandomRotate90, MedianBlur, ShiftScaleRotate, OpticalDistortion, GridDistortion, and HueSaturationValue

#### Training

Training for the first epoch was conducted with Automatic Mixed Precision (AMP) enabled to enhance computational efficiency on our hardware. The SupCon loss function had a temperature setting of 0.1. The Exponential Moving Average (EMA) with a decay rate of 0.4 per epoch was utilized to stabilize training outcomes. The primary training metric was set to precision at one, with Stochastic Gradient Descent (SGD) as the optimizer at a learning rate of 0.003. The learning rate was modulated using a CosineAnnealingLR scheduler, which reduced the rate following a cosine curve from its initial value to 0.0003 over the training period.

## Results

In the first stage, after training for 100 epochs, the training loss decreased from 6.464 ± 0.012 to 5.156 ± 0.012, and the encoder validation metrics showed a consistent increase in precision at one, from its initial value of 0.564 ± 0.056 to an average of 0.94 ± 0.012 by the end of the training period. In the second stage the training loss significantly decreased from 0.217 ± 0.014, to 0.141 ± 0.01, whereas the validation loss decreased from 0.179 ± 0.037 to 0.198 ± 0.045.

## Notes

https://github.com/agporto/BioEncoder

https://zenodo.org/records/10909614

### Competing Interest Statement

The authors have declared no competing interest.

